# Cryo-EM structure of amyloid fibrils formed by the entire low complexity domain of TDP-43

**DOI:** 10.1101/2020.12.14.422689

**Authors:** Qiuye Li, W. Michael Babinchak, Witold K. Surewicz

## Abstract

Amyotrophic lateral sclerosis and several other neurodegenerative diseases are associated with brain deposits of TDP-43 aggregates. Cryo-EM structure of amyloid formed from the entire TDP-43 low complexity domain reveals single protofilament fibrils containing a large (138-residue), tightly packed core with structural features that differ from those previously found for fibrils formed from short protein fragments. The atomic model provides insight into potential structural perturbations caused by phosphorylation and disease-related mutations.

---

Amyloid-like aggregates formed by 25-35 kDa C-terminal fragments of TDP-43 are a pathological hallmark of amyotrophic lateral sclerosis (ALS), frontotemporal lobar degeneration (FTLD), and several other neurodegenerative diseases^1,2^. Aggregation of TDP-43 is largely driven by the low complexity domain (LCD) of the protein^3,4^. In contrast to recent progress in high-resolution structural studies of other amyloids^5–17^, however, similar studies with TDP-43 are limited to fibrils formed by relatively short fragments of the protein^18^. To bridge this critical gap, here we report a near-atomic resolution cryo-electron microscopic (cryo-EM) structure of fibrils formed by the entire LCD domain of TDP-43.

Cryo-EM images of TDP-43 LCD fibrils revealed two types of morphologies, one of them showing a helical twist and the second one lacking such a twist (Extended Data Fig. 1a). Helical reconstruction of twisted fibrils (all of which share the same fold) allowed us to determine a density map with a nominal resolution of 3.2 Å (Extended Data Table 1 and Extended Data Fig. 1b). Each twisted fibril consists of a single left-handed protofilament in which subunits stack along the fibril axis with a helical rise of 4.73 Å and a helical twist of −1.66° (Fig. 1a). A near-atomic-resolution structural model could be unambiguously built for the fibril core, which maps to residues 276-414 of TDP-43 (Fig. 1b and Extended Data Table 1). To the best of our knowledge, the 138-residue core region of TDP-43 LCD fibrils represents the largest amyloid core reported to date.

**Fig. 1.**
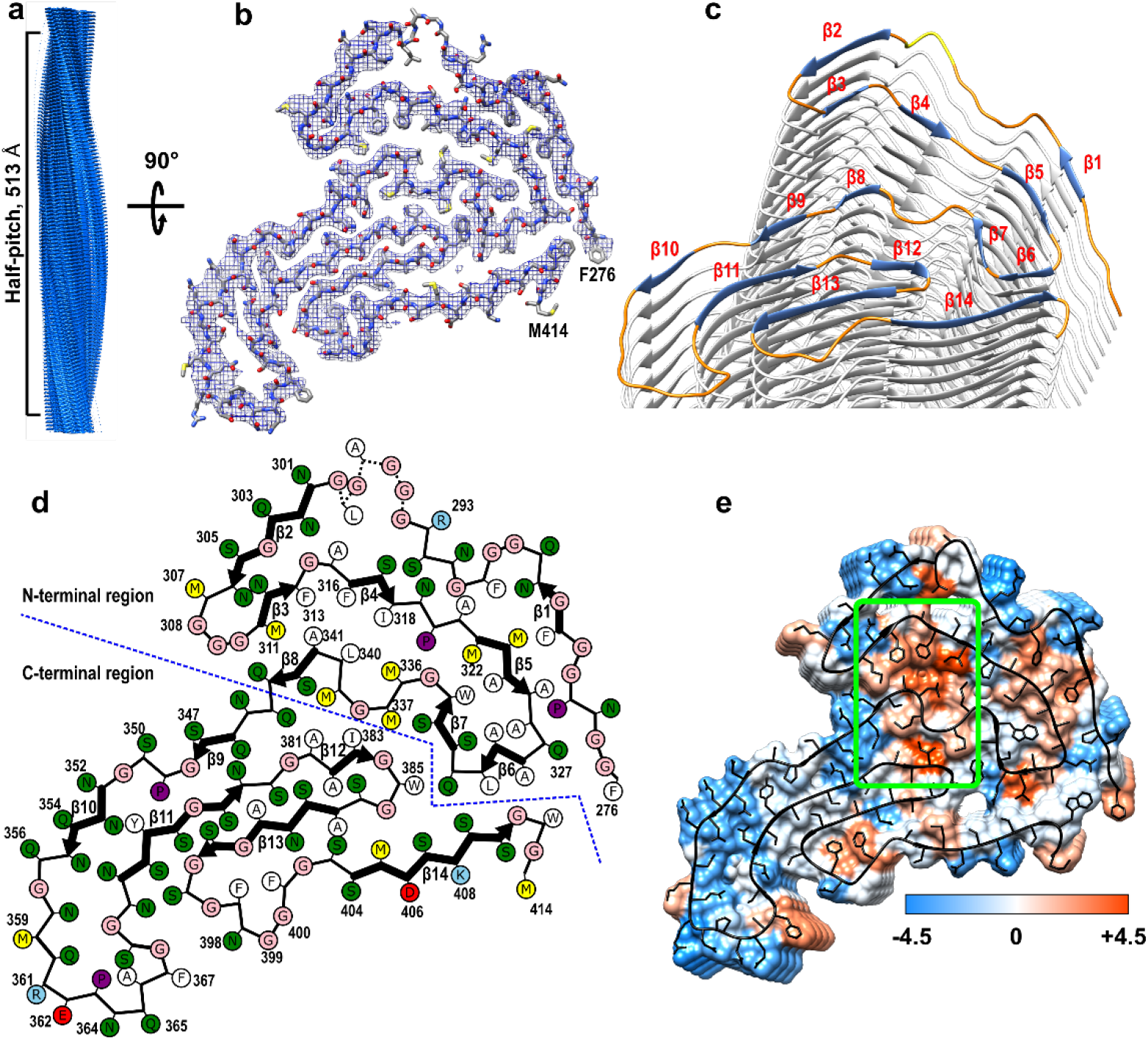
Cryo-EM structure of TDP-43 LCD fibrils. **a,** Cryo-EM density map showing a left-handed helix with a half-pitch of 513 Å. Repeating densities representing β-strands along the z-axis with a 4.73 Å interval indicate a parallel in-register β-sheet architecture. **b,** The atomic model and superimposed cryo-EM map of one cross-sectional layer of each fibril (top view). **c,** Tilted cross-section of a TDP-43 LCD fibril. β-strands in the top subunit are shown in blue, highly ordered turns in orange, and the less ordered region in yellow. **d**, Schematic representation of one cross-sectional layer of the amyloid core, with β-strands shown as thicker arrows and the less ordered region (residues 295-299) marked as dotted lines. **e**, Hydrophobicity of the fibril cross-section, with hydrophobicity levels colored according to Kyte-Doolittle^20^. A major hydrophobic core (green box) is made up of the 311-318, 336-341, and 381-383 segments.

This core consists of 14 β-strands linked by relatively rigid turns and loops (Figure 1c and d). Each protein subunit can be divided into two regions that differ with regard to an overall amino acid composition as well as the backbone geometry (Fig. 1d and Extended Data Fig. 2). The C-terminal region (residues 344-414, strands β9-β14) is characterized by a small proportion of hydrophobic amino acids. Relatively long β-strands pack mostly via polar interfaces and many residues within strands β9, β10, β11 and β13 are engaged in steric zipper interactions (Fig. 1d and Extended Data Fig. 2a). The stacking of subunits within this region is largely maintained by a network of intermolecular backbone H-bonds between β-strands, with additional stabilization through H-bonds between numerous stacked Gln and Asn side chains (Extended Data Fig. 3). This C-terminal region is largely planar, with most β-strands packed through interactions within the same subunit (Extended Data Fig. 2b).

By contrast, the N-terminal region of LCD (residues 276-343, strands β1-β8) is rich in hydrophobic amino acids and contains mostly short β-strands that are not involved in steric zipper interactions. Instead, strands and turns pack against each other in all directions, allowing non-polar side chains to be tightly packed and buried, with a hydrophobic interface between the 311-318 and 336-341 segments (Fig. 1d and e). Side chains on the opposite side of the 336-341 segment form another hydrophobic interface with residues 381-383 within the LCD C-terminal region. Altogether, these three segments make up a large hydrophobic core. A second, smaller hydrophobic core involves residues within the Ala- and Met-rich 321-330 segment that pack against Phe283, Phe289, and Trp334 (Fig. 1e and Extended Data Fig. 2a). To facilitate such a tight packing of hydrophobic residues, the backbone in this region of each subunit makes numerous turns within the x-y plane while also extending along the z-axis over a distance of ~22.4 Å (Extended Data Fig. 2b). As a result, each subunit (*i*) not only interacts with the layer directly above (*i+1*) and below (*i-1*), but also with layers up to (*i+3*) and (*i-3*), as illustrated in Fig. 2a for the 311-327 segment. Furthermore, some of the interactions are between stacks of hydrophobic residues which are arranged in a staggered fashion (Fig. 2b-c). Thus, even though only ~40% of residues are involved in intermolecular H-bonds within the cross-β motif, fibrils are likely further stabilized through the interlayer interactions between side chains.

**Fig. 2.**
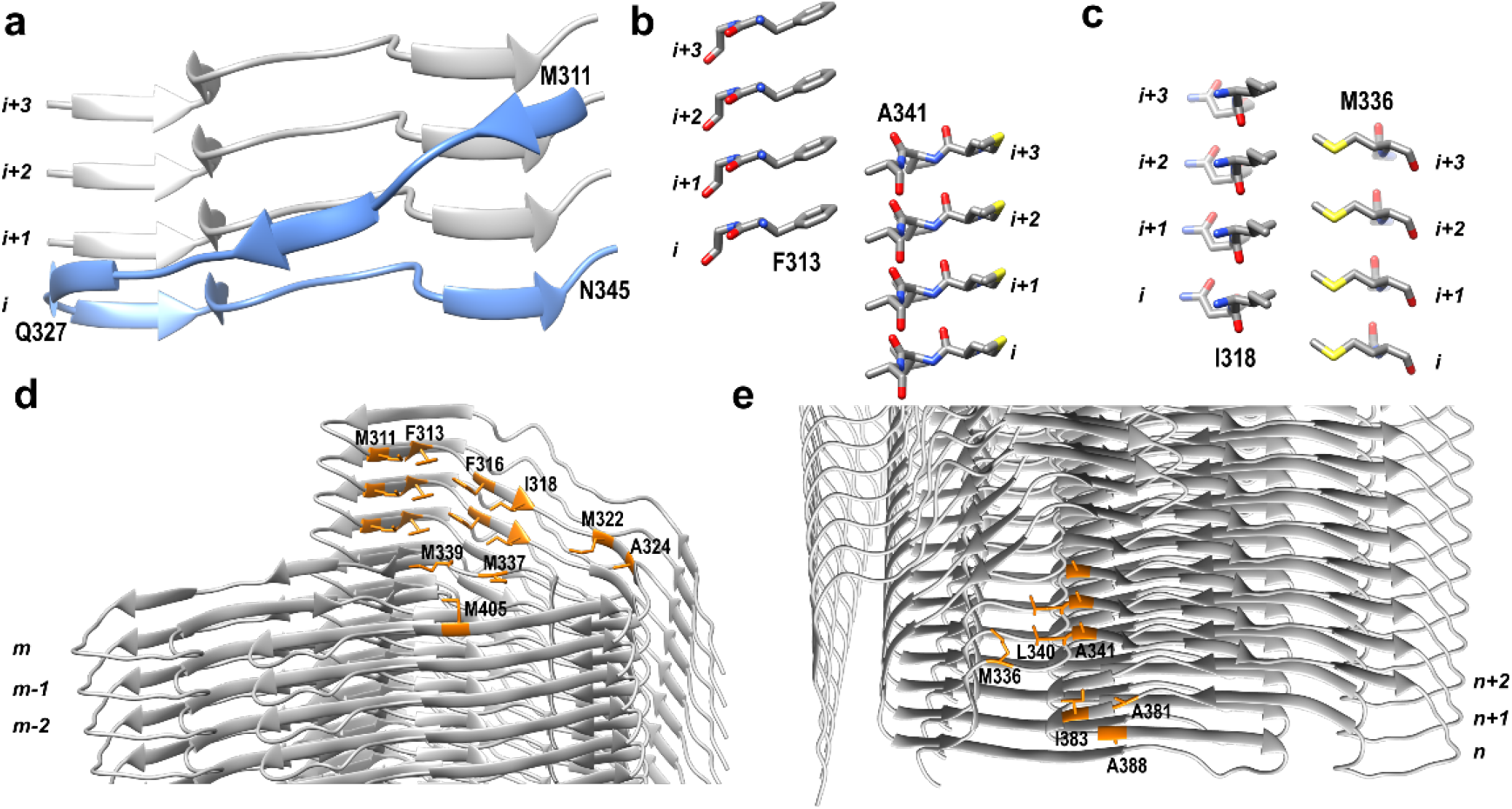
Structural features of the N-terminal region of TDP-43 LCD amyloid core. **a**, Side view of the 311-327 segment extending along the fibril axis. This segment in subunit *i* interacts with the 327-345 segment in subunits *i+1*, *i+2* and *i+3*. **b and c**, Side view of interlayer interactions between F313 and A341 (**b**) and I318 and M336 (**c**). Layers of side chains are packed in a staggered fashion, resulting in a very compact hydrophobic core. **d and e**, Solvent accessible hydrophobic amino acids (orange) at the top (**d**) and bottom (**e**) end of the fibril. Fifteen hydrophobic side chains from different layers [M331 (*m*, *m-1*, *m-2*), F313 (*m*, *m-1*, *m-2*), F316 (*m*, *m-1*), I318 (*m*, *m-1*), M322 (*m*), A324 (*m*), M337 (*m*), M339 (*m*), and M405 (*m*)] are exposed to water at the top end. Nine hydrophobic side chains [A341 (*n*, *n+1*, *n+2*), L340 (*n*, *n+1*), M336 (*n*), A381 (*n*), I383 (*n*), and A388 (*n*)] are exposed at the bottom end.

The non-planar backbone conformation of TDP-43 LCD subunits (which is not unusual among amyloids^6,11,15^) results in rugged surfaces of fibril ends, with fifteen hydrophobic residues from the top three subunits (*m*, *m-1*, *m-2*) and nine hydrophobic residues from the bottom three subunits (*n*, *n+1*, *n+2*) exposed to water (Fig. 2d and e and Extended Data Fig. 4). Such highly hydrophobic, rugged surfaces of fibril ends may be especially conducive to the recruitment of monomers and their templated conversion into the amyloid conformation. Furthermore, distinct surfaces at opposite ends could result in fibril polarity, with different elongation rates at each end.

A recent study reported cryo-EM structures of fibrils formed by two relatively short fragments of TDP-43 LCD^18^. The first fragment (residues 311-360) formed polymorphic amyloid structures with a common motif described as a dagger fold, in which residues from Phe313 to Ala341 form tight hydrophobic interaction. A sharp (~160°) kink at Gln327 defines the tip of the dagger (Fig. 3a and Extended Data Fig. 5a). The fold adopted by this segment in our structure of fibrils formed by the entire LCD is substantially different, with no sharp kink at Gln 327. Instead, there is a ~90° turn at this residue, followed by another ~90° turn at Gln331, such that the overall fold shows little resemblance to the dagger motif (Fig. 3a and Extended Data Fig. 5b). Fibrils formed by the second fragment (residues 286-331 with A315E mutation) consisted of four protofilaments that each contain another common motif characterized as R-shaped fold Fig. 3b and Extended Data Fig. 5c). It was proposed that a similar fold (stabilized by hydrophobic interactions between Ala315, Ala297 and Phe313) would be adopted by this segment without the mutation^18^. Again, the fold within this region of fibrils formed by the entire LCD is quite different: Ala297, Phe313, and Ala315 are not in close contact, and the local conformation is dictated by interactions with other parts of the molecule (Fig. 3b and Extended Data Fig. 5d).

**Fig. 3.**
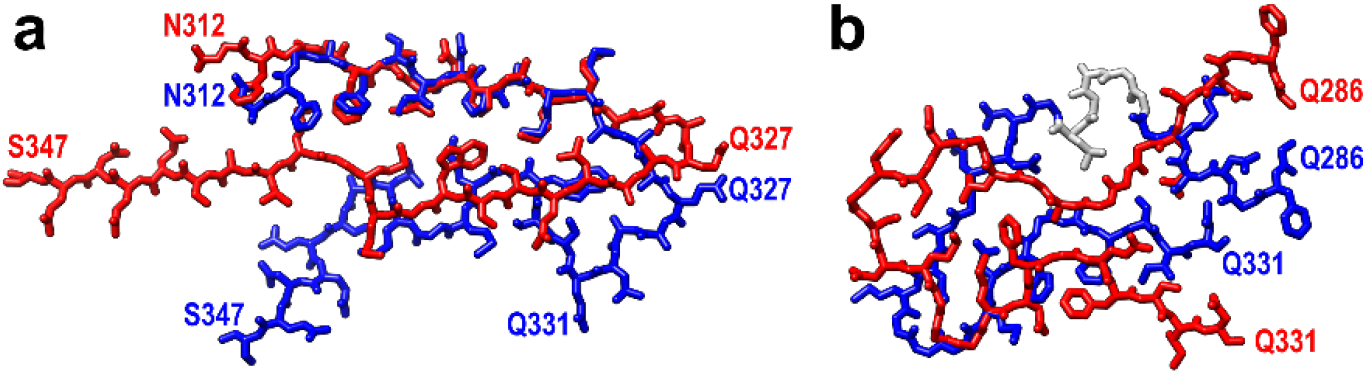
Comparison of structural motifs in fibrils formed by relatively short fragments of TDP-43 LCD^18^ with those in the same regions of fibrils formed by the entire LCD. **a**, Representative dagger-shaped fold of the 312-347 region in fibrils formed by the 311-360 fragment (red, PDB 6N37) compared to the same region in TDP-43 LCD fibrils (blue). **b**, The R-shaped fold in the inner protofilament of fibrils formed by the 286-331 fragment with A315E mutation (red, inner protofilament in PDB 6N3C) compared to the structure of the relevant region within TDP-43 LCD fibrils (blue; a less-defined region shown in grey).

Over thirty point mutations within the TDP-43 LCD are associated with ALS and FTLD^1,2^. The mechanisms by which these mutations facilitate disease are poorly understood. Mapping the pathogenic mutations on the structure of wild-type TDP-43 LCD fibrils revealed that ~50% of them are not compatible with this structure due to steric clashes within tightly packed segments of the protein, introduction of charges into the dehydrated fibril interior, or both (Extended Data Fig. 6a). Thus, these mutations will likely result in substantially different fibril structures, and this may affect the disease phenotype.

One of the features of ALS and FTLD pathology is phosphorylation of Ser residues within the C-terminal part of TDP-43 LCD^2,19^. Most of these phosphorylation sites are buried in the interior of the fibril structure that we determined for the non-phosphorylated protein (Extended Data Fig. 6b). Thus, phosphorylation of each of these individual residues will likely affect the structure of fibrillar aggregates, resulting in a large, phosphorylation site-dependent structural polymorphism. This, in turn, could further influence the disease phenotype.

## Methods

### Protein Expression and Purification

TDP-43 LCD with an N-terminal His-tag was expressed and purified as described previously^4^. In short, protein was expressed overnight using Rosetta™ (DE3) pLysS competent cells (MilliporeSigma) after induction with 1 mM isopropyl β-D-1-thiogalactopyranoside. Cells were collected by centrifugation, lysed by sonication, and purified over Ni-charged nitrilotriacetic acid column using 4-5 column volume washes with a pH 8 buffer containing 8 M urea followed by elution with 250 mM imidazole. Protein was concentrated and purified by HPLC using a C_4_ column with acetonitrile gradient in water containing 0.05% trifluoroacetic acid. Protein purity (> 95%) was confirmed by gel electrophoresis. Pure protein was flash frozen and lyophilized for later use.

### Fibril Formation

Lyophilized protein was dissolved in Milli-Q purified H_2_O and passed through an Amicon Ultra centrifugal filter with 100 kDa molecular weight cut-off to remove preformed aggregates^4^. Fibrils were formed at a protein concentration of 30 μM in 20 mM sodium acetate buffer, pH 4.0. The sample was placed on rotation (~ 6-8 rpm) at 37°C. When analyzed by atomic force microscopy, the resulting fibrils displayed two different morphologies, one twisted and one lacking such a twist. In an attempt to select one preferential morphology, four rounds of sequential seeding reactions were then performed by adding preformed, sonicated fibrils (10% w/w) to freshly prepared, non-aggregated protein under the same buffer conditions. Despite these efforts, akin to the first-round fibrils, the final sample used for cryo-EM studies also contained two types of fibril morphologies (Extended Data Fig. 1).

### CryoEM

200 mesh lacey carbon grids (Ted Pella) were first coated with 0.1 mg/ml graphene oxide and then with 0.1% poly-lysine as described previously^21,22^. 3 μl of TDP-43 LCD fibril suspension (30 μM) was applied to the coated grid, blotted for 6 s, and plunge-frozen in liquid ethane using a Vitrobot Mark IV (ThermoFisher Scientific). Movies were collected on a Titan Krios G3i microscope (ThermoFisher Scientific) equipped with a BioQuantum K3 camera (Gatan, Inc), with 0.414 Å/pixel at Super Resolution mode, 42 e^−^/Å^2^ total dose and 60 total frames. A total of 6,589 micrographs were automatically collected using SerialEM^23^ with 6 shots per position. Beam image shift was applied, and defocus range was between −0.8 and 1.5 μm,

### Data processing

Movies were corrected for drifting and binned by a factor of 2 using MotionCor2^24^. Contrast Transfer Functions (CTF) were estimated by Gctf^25^. All further processing was carried out using RELION 3.1^26–28^. Fibrils with apparent helical twists were manually picked and a total of 294,168 segments were extracted using an overlap of 97% between neighboring segments and a box size of 512 pixels. Segments were first subjected to several rounds of reference-free 2D classification using T=8 and K=100 to remove poorly defined classes, resulting in 65,075 segments contributing to clear 2D averages. These segments were then used for subsequent 3D classification employing an initial model of a featureless cylinder generated by *relion_helix_toolbox*^27^. The initial helical rise (4.73 Å) was calculated from the 2D class layer line profile, and the initial helical twist (−1.64°) was calculated from the crossover distance. The handedness of the helix was determined by atomic force microscopy images. The tilt of all segments were kept at 90° throughout the 3D processing. Two rounds of 3D classification were performed using K=3 and T=4, resulting in 11,026 segments that contributed to a high-resolution reconstruction. Additional rounds of 3D classification were performed on these segments using a single class and increasing T value (4, 8, 20, 40, and 100) with local optimization of helical twist and rise. In the last round of 3D classification, β-strands were well separated and large side chains could be resolved. The model and data were then used for high-resolution gold-standard 3D refinement. Iterative Bayesian polishing^28^ and CTF refinement^29^ were performed to further improve the resolution. The overall resolution was calculated to be 3.2 Å from Fourier shell correlations at 0.143 between two independently refined half-maps. Refined helical symmetry (twist = −1.66°, rise= 4.73 Å) was imposed on the post-processed map for further mode building.

### Model building

An initial model was built in *Coot*^30^ using large side chains of ^343^QQNQ^346^ segment as a guide. Five chains were built at the central region of the density map, which covered all types of intermolecular contact within the map. The model was then subjected to iterative real-space refinements in PHENIX^31,32^. At later stages, segments favoring β-strand conformation were identified and the direction of backbone oxygen and nitrogen atoms were adjusted manually to facilitate hydrogen bonding in β-sheets. Such restraints were also implemented in the subsequent refinements. After real-space refinement, side-chain orientations were manually adjusted to ensure energy-favored geometry. The final model was validated using the comprehensive validation method in PHENIX^33,34^.

The density of the TDP-43 LCD segment 295-299 was weaker than that of the rest of the molecule, possibly due to the less ordered structure of this segment. In this region, we displayed the map at a low contour level and built the model manually. After automatic refinement in PHENIX^31,32^, this region showed no significant clashes or Ramachandran outliers. It has been reported that such less-defined turns or loops of an amyloid core may adopt more than one conformation^35^. In our study, due to the relatively low number of particles, we were not able to classify and determine different structures in this less-defined region. Thus, the structure of this segment in our refined model may represent the average of multiple conformations.

When assessing structural compatibility of disease-related mutations and phosphorylation at individual Ser residues with our structural model determined for wild-type, non-phosphorylated protein, the *Reduce*^36^ and *Probe*^37^ programs were used to test for steric clashes in the absence of the backbone movement.

## Acknowledgements

This work was supported by NIH grants AG061797 and GM094357. We thank Kunpeng Li for help with acquisition of cryo-EM data, Gunnar F. Schröder and Xinghong Dai for discussions regarding data processing and model building, respectively, and Sudha Chakrapani for comments on the manuscript.

## Author contributions

Q.L. and W.K.S. designed the study. Q.L. collected EM data, performed image processing and model building. W.M.B. purified the protein and prepared fibrils. Q.L. and W.K.S. wrote the manuscript.

## Competing interests

The authors declare no competing interests.

## EXTENDED DATA

**Extended Data Fig. 1.**
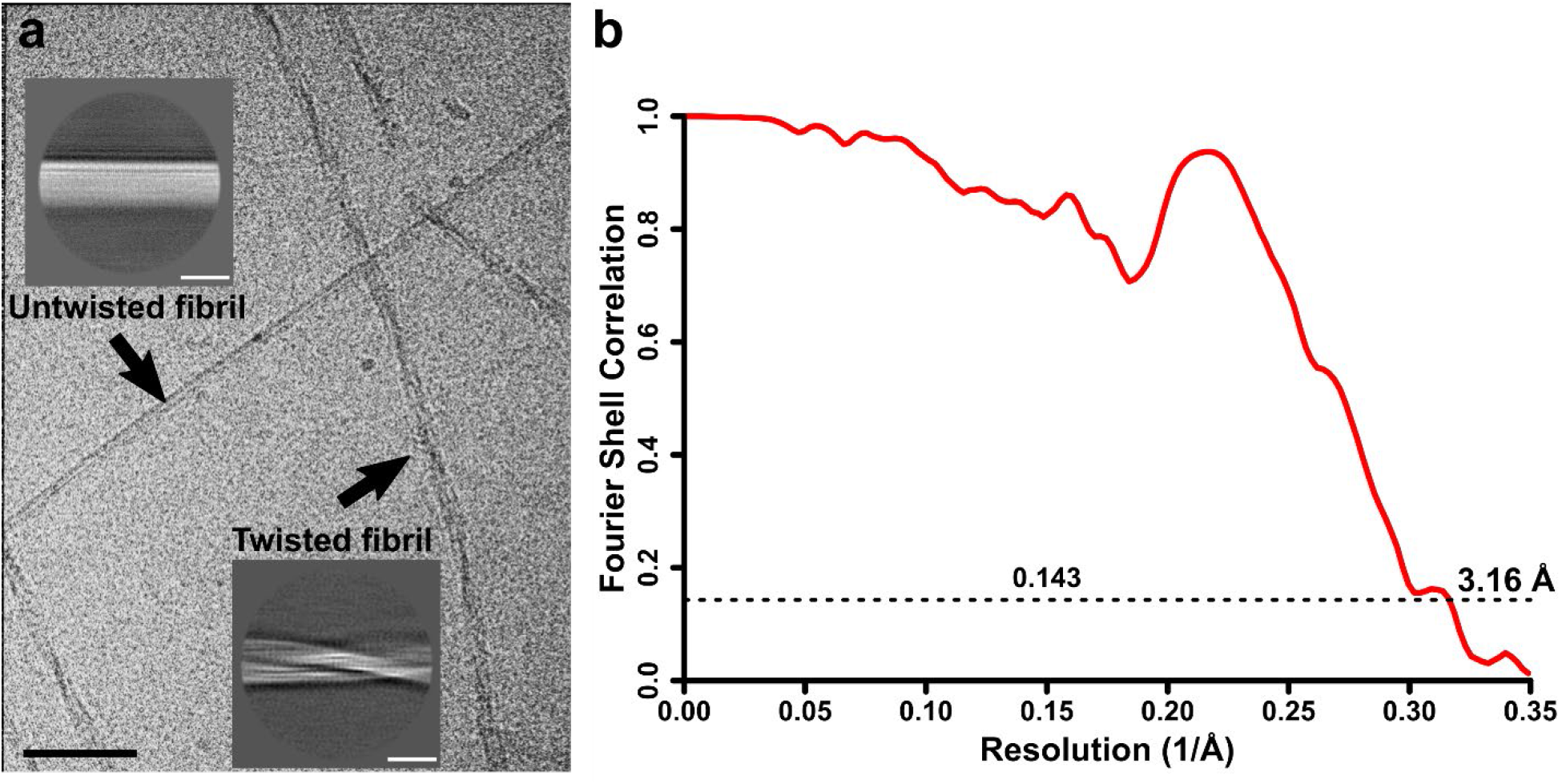
Cryo-EM data processing. **a,** Two different types of morphologies found for TDP-43 LCD fibrils. Cryo-EM images show both twisted and untwisted fibrils; scale bar (black) corresponds to 50 nm. 2D class averages of each fibril type are shown as insets; scale bars (white) correspond to 10 nm. **b**, Fourier shell correlation curve between the two independently refined half-maps.

**Extended Data Fig. 2.**
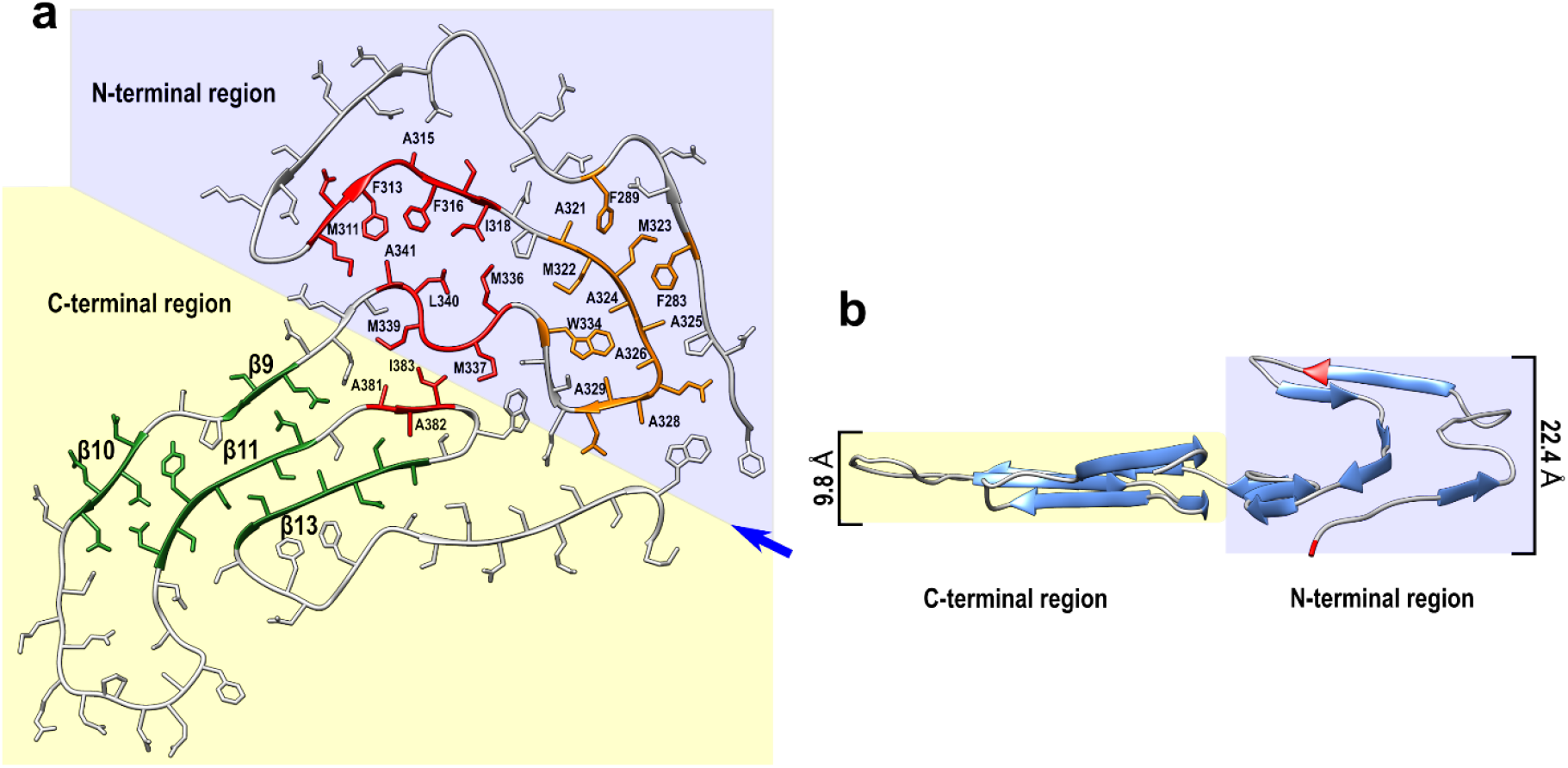
Differences between the N- and C-terminal regions of the amyloid core of TDP-43 LCD fibrils. **a**, Top view of one subunit within the fibril. Large (red) and smaller (orange) hydrophobic cores are present within the N-terminal part, stabilizing this region. The C-terminal region is stabilized largely by steric-zipper interactions involving side chains within strands β9, β10, β11, and β13 (green). **b**, Side view of one subunit within the fibril (with the view angle indicated by the blue arrow in panel a). In contrast to a largely planar C-terminal region, the N-terminal region extends along the fibril axis over the distance of 22.4 Å. The lowest (Phe276) and the highest (Asn306) points are marked in red.

**Extended Data Fig. 3.**
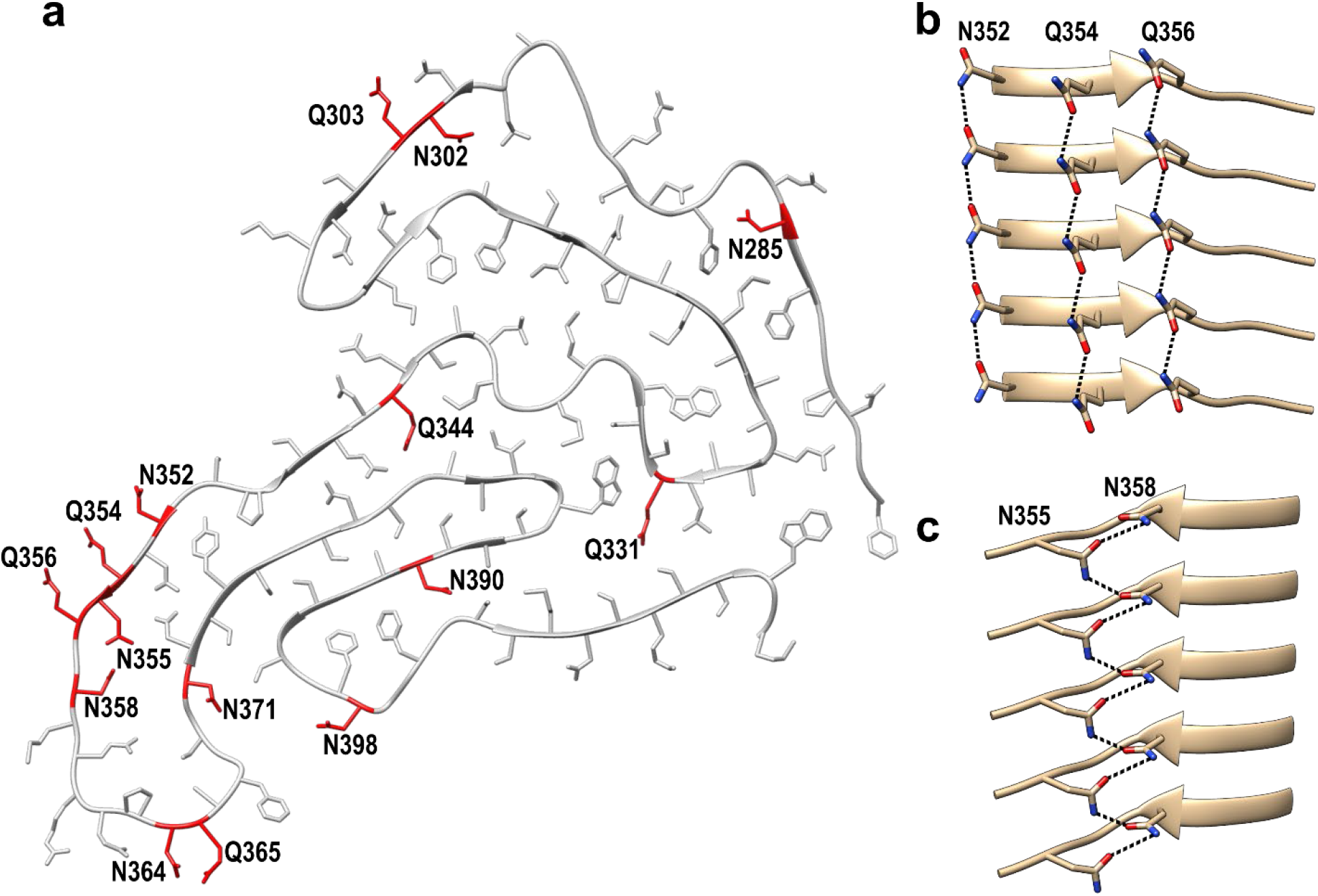
Illustration of stabilizing interlayer hydrogen bonds between stacked Asn and Gln side chains. **a**, Top view showing Asn and Gln residues (marked in red) involved in interlayer hydrogen bonding. **b**, Asn/Gln ladders of Asn352, Gln354, and Gln356 with hydrogen bonds (black dashed lines) between the same residues in adjacent subunits. **c**, Ladders of Asn355 and Asn358 in which hydrogen bonds are between different residues in adjacent subunits.

**Extended Data Fig. 4.**
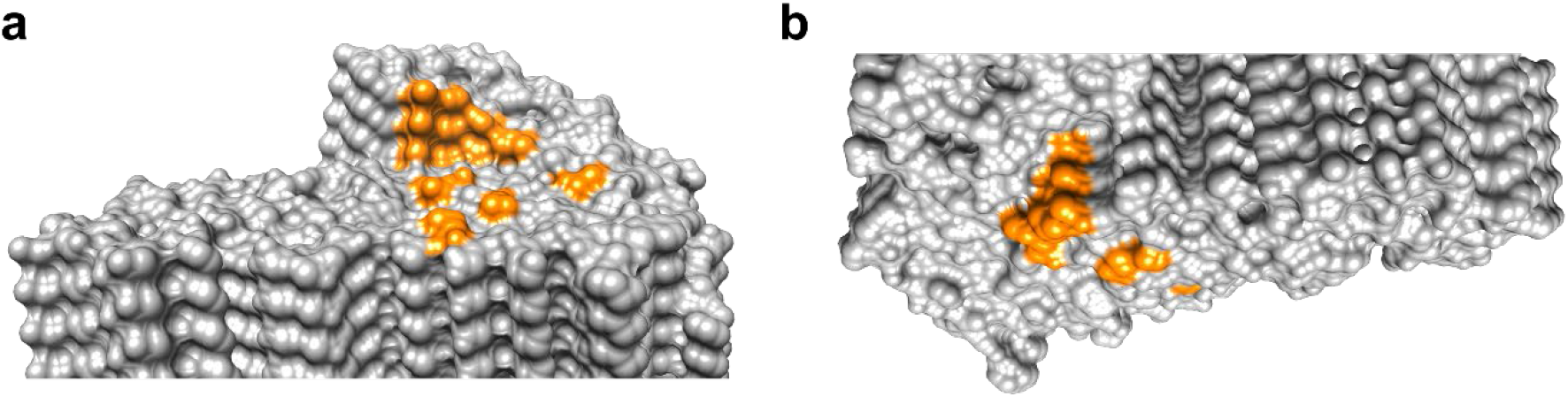
Surface representation of the top (a) and bottom (b) ends of TDP-43 LCD fibril. Solvent accessible hydrophobic residues are marked in orange.

**Extended Data Fig. 5.**
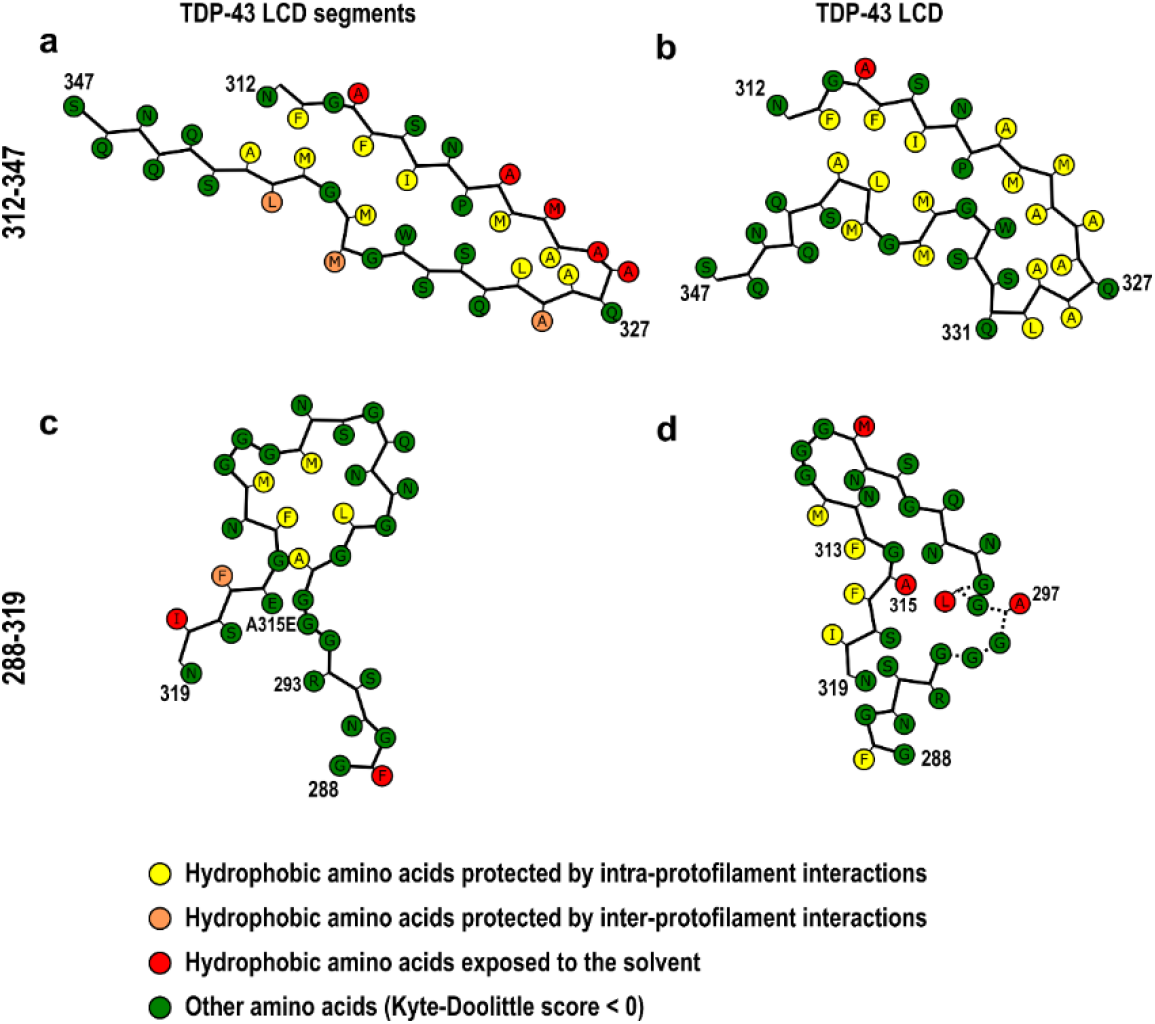
Comparison of structural motifs in fibrils generated from relatively short fragments of TDP-43 LCD^18^ with those in the same regions of fibrils generated from the entire LCD. **a**, Representative structure of fibrils formed by the 311-360 fragment (PDB 6N37). Almost half of hydrophobic residues within the dagger motif (residues 312-347) are either exposed to water or protected by intermolecular interactions with another protofilament (only one protofilament is shown here). **b**, The structure of this region within a much larger amyloid core of fibrils formed by the entire LCD is substantially different. This region is buried in the interior of a single protofilament, with almost all hydrophobic residues involved in intra-protofilament interactions and protected from water (see also Fig. 1d). **c**, The structure of the inner protofilament in fibrils formed by the 286-331 fragment with A315E mutation showing an R-shaped fold within the 288-319 segment, with a salt bridge between Arg293 and Asp315 (PDB 6N3C). A similar R-shaped fold was proposed to be adopted by fibrils formed by this fragment in the absence of any mutation, with the structure stabilized by hydrophobic interactions between Ala297, Phe313, and Ala315. **d**, The structure within the 288-319 region of fibrils formed from the entire LCD is quite different.

**Extended Data Fig. 6.**
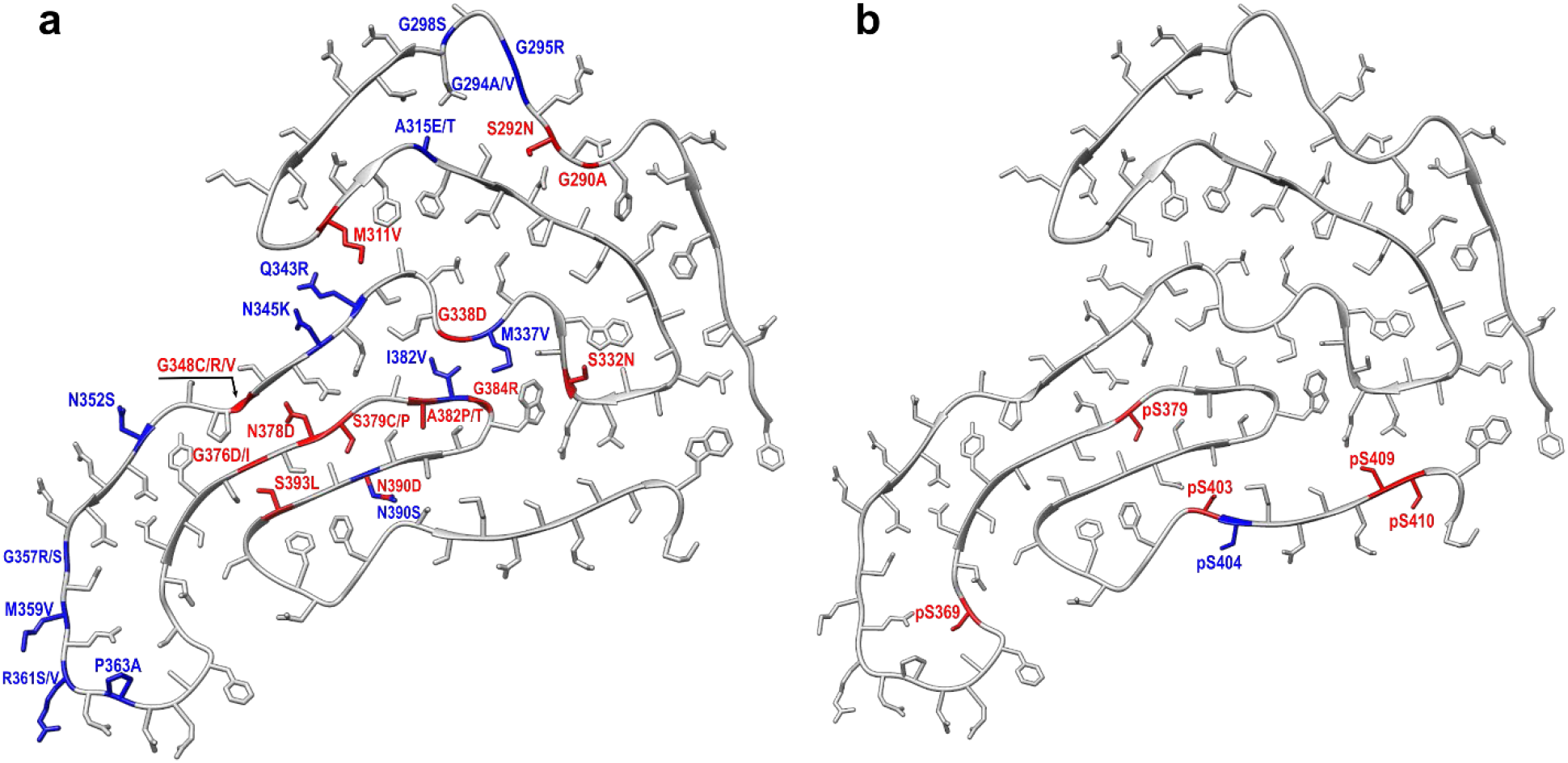
Disease-related point mutations and phosphorylation sites mapped on the structure of one subunit of fibrils formed from the wild-type, non-phosphorylated TDP-43 LCD. **a**, Mutations that are compatible with this structure are labeled in blue. The remaining mutations (labeled in red) are not compatible with the structure determined for wild-type protein fibrils due to steric clashes within tightly packed segments of the protein (G290A, S292N, M311V, S332N, G348C/V, G376I, S379C/P, A382P/T, S393L), introduction of charges into the dehydrated fibril interior (G338D, G348R, G376D, N378D, N390D), or both (G338D, G348R, G376D). **b**, Phosphorylation sites exposed on the surface and those buried inside the structure determined for wild-type TDP-43 LCD fibrils are labeled in blue and red, respectively.

**Extended Data Table 1.** Cryo-EM data collection, refinement, and validation statistics

|  |  |
| --- | --- |
| **Data collection and processing** |  |
| Magnification | × 105,000 |
| Voltage (kV) | 300 |
| Electron dose (e<sup>-</sup>/Å<sup>2</sup>) | 42 |
| Defocus range (μm) | -0.8 to -1.5 |
| Pixel size (Å) | 0.828 |
| Symmetry imposed | C1 |
| Initial particles images (no.) | 294,168 |
| Final particle images (no.) | 11,026 |
| Map Resolution (Å) | 3.16 |
| FSC threshold | 0.143 |
| **Refinement** |  |
| Initial model used | De novo |
| Map sharpening B factor (Å<sup>2</sup>) | -29.83 |
| Model composition |  |
| Non-hydrogen atoms | 4,695 |
| Protein residues | 695 |
| B factors (Å<sup>2</sup>) |  |
| protein | 94.85 |
| R.M.S.D |  |
| Bond lengths (Å) | 0.006 |
| Bond angles (°) | 1.170 |
| MolProbity score | 2.07 |
| Clashscore | 12.59 |
| Poor rotamers (%) | 0 |
| Ramachandran plot |  |
| Favored (%) | 92.70 |
| Allowed (%) | 7.30 |
| Disallowed (%) | 0 |

## Notes

### Competing Interest Statement

The authors have declared no competing interest.

